# Novel method for the extraction of DNA from Lamniformes tooth and denticle enamel Suitable for PCR

**DOI:** 10.1101/2021.01.18.427093

**Authors:** Bryan Swig, Ralph S. Collier

**Affiliations:** California Lutheran University, 60 West Olsen Road, #3700Thousand Oaks, California 91360; Shark Research Committee, P. O. Box 3492Chatsworth, California 91313

**Author notes:** Corresponding author (BS).

**Keywords:** DNA, Extraction, Tooth, Denticle, Enamel

## Abstract

Extraction of high-quality genomic DNA from Lamniformes tooth fragment and dermal denticle enamel is discussed as a method of identifying individual sharks. We describe a procedure that permits isolation of genomic DNA of satisfactory size and quality for PCR analysis, as well as for most routine cloning applications. This method should allow for the non-invasive collection of genomic samples from Lamniformes.

## Introduction

Historically, tooth fragments have been deposited by Lamniformes in both inanimate and animate objects worldwide. Fragments, usually from a white shark, Carcharodon carcharias, removed from inanimate objects, prey, and shark/human interactions along the Pacific Coast of North America, both fatal and non-fatal, have provided only insights into species identification and not a specific individual shark. Most notable of these prior cases involving the recovery of tooth fragments are described by; Ames & Morejohn 1966; Coppleson 1954; Baldridge 1974; Collier 1964, 2003; Collier et. al.1996; Davies 1964; Follett 1974; Miller & Collier 1981; Wallett 1978; and Tinker et. al. 2015. With the ever expanding DNA data base of elasmobranchs globally a method for obtaining profiles from minute amounts of teeth seemed appropriate. Further, obtaining specific identification of the sharks involved in these events can provide insights into their movements for future conservation suggestions and public awareness as was the case with conservations efforts for African elephants (Mailand, C. & S. K. Wasser 2007).

While blood and muscle tissue biopsy’s represent the most commonly used sources of DNA utilized in genetic studies of Lamniformes (Keeney et. al. 2005; Feldheim et. al. 2001; Taguchi ed. al. 2013) an alternative methodology was required for tooth fragments and dermal denticles. While the information provided from these common tissue and blood samples are invaluable, obtaining these traditional samples can be both difficult and potentially harmful to the organism as observed in the injuries sustained from SPOT tag deployments (Jewell, O.J.D., et. al 2011; Hammerschlag, N R., et.al. 2011; R. Warner, pers. comm.). Therefore, non-invasive sampling is a very attractive alternative, allowing for genetic analysis without having to catch or handle the specimen (Wasko et. al. 2003). DNA can be obtained from a variety of samples including enamel from scales and teeth. In the present paper, a DNA extraction method is described as template in polymerase chain reaction (PCR) experiments.

## Materials and Methods

### Sample Collection and Processing

For this study a single *Isurus oxyrinchus* was utilized to collect tissue, denticle, and tooth enamel samples. This sample was donated to us by a local fisherman. California Lutheran Universities IACUC committee provided us with a waiver as no live organisms were used in this study.

For each tissue sample, a 1 to 1.5 g plug of dorsal white muscle was taken 5 to 10cm ventral to the base of the first dorsal fin and 2 to 5 cm below in what is typically part of the carcass dressed out for human consumption. Muscle plugs were removed using clean, stainless steel instruments and were placed in sterile microcentrafuge tube.

The jaw of the sample was removed and dried at room temperature in a chemical fume hood for 1 week. Following drying the teeth were removed from the jaw by cutting them free with a jewelers saw. Teeth were then placed into microcentrafuge tubes.

All samples used in this experiment were collected simultaneously, and immediately frozen at −20°C. Samples stayed frozen at −20°C until extraction took place.

## DNA Extraction from Tooth Fragments

Teeth were fragmented with extreme care to avoid the dentine and pulp of the tooth, thus leaving samples that were comprised of only enamel. Tool, cutting parts, surfaces, and vice were washed with 70 % ethanol and 10 % sodium hydroxide. Samples were placed into 1.5 ml microcentrifuge tubes. Tooth samples (averaging 0.20 g) were initially washed, and subsequently crushed to powder on a Spex 6770 freezer mill (Spex SamplePrep). Pulverized samples were placed in 1.5 ml microcentrifuge tubes with 1 ml of EDTA pH 8.0, and incubated at 37°C for 24 hours with occasional shaking. Samples were then spun at 9000g for 10 minutes, the resulting supernatant was discarded and the pellet was re-suspended in 1 ml of EDTA pH 8.0, and incubated at 37°C for 24 hours with occasional shaking. Following the second incubation period samples were again then spun at 9000g for 10 minutes, the resulting supernatant was discarded and the pellet was then washed in 1 ml of nuclease free water Samples were again spun at 9000g for 10 minutes, the supernatant was discarded and the pellet was re-suspended in 1 ml of nuclease free water then spun at 9000g for 10 minutes, the supernatant was discarded. The pellet was then re-suspended in 360 μl of ATL^*^ lyse buffer and 40 μl of Proteinase K^*^ was added and the samples were maintained at 56°C for at least 10 h. 400 μl AL^*^ buffer was then added to each sample, then mixed. Samples were then maintained at 70°C for 10 minutes. Following the 10 minute incubation 400 μl of 100% EtOH was added to each sample and mixed. 675 μl of each sample was then placed into a spin column and spun at 6000g for 1 min, the remaining sampled was then added to the spin column and spun at 6000g for 1 min. 500 μl of AW1^*^ buffer was added to the column and spun at 6000g for 1 min. 500 μl of AW2^*^ buffer was added to the column and spun at 15000g for 3 min. The column was then moved into a 2ml screw cap tube and 100 μl of AE^*^ buffer was added. Samples were then maintained at 21°C for 10 minutes, following the incubation period samples were spun at 9000 g for 1 minute. 100 μl of AE buffer was again added and, samples were then maintained at 21°C for 10 minutes, following the incubation period samples were spun at 9000 g for 1 minute resulting in a yield of 200 μl eluate, that can be stored at −20°C until analyses could occur. (^*^ ATL, AL, AE, AW1, AW2 buffers and Proteinase K are from Qiagen, Valencia, CA, USA)

## DNA Extraction from Tissue Samples

DNA extraction from tissue samples followed the methods and procedures outlined in the Quiagen DNeasy Blood and Tissue hand book (Qiagen, Valencia, CA, USA). DNA extraction from tooth samples followed the procedure outlined above. After extraction samples were frozen at −20°C until analyses could occur.

## Concentration and Purity Determination

A quantitative spectrophotometric assay of DNA was performed using a NanoDrop UV-visible spectrophotometer (Thermo Scientific, Canoga Park, CA, USA). Absor-bance was measured at wavelengths of 260 and 280 (A_260_ and A_280_, respectively) nm. The absorbance quotient (OD_260_/OD_280_) provides an estimate of DNA purity. An absorbance quotient value of 1.8 < ratio (R) <2.0 was considered to be good, purified DNA. A ratio of <1.8 is indicative of protein contamination, where as a ratio of >2.0 indicates RNA contamination.

## Results and Discussion

The amount of DNA recovered from a tooth fragment and denticle varied between extracts. The average yield from a fragment was 22.38 ug/ul, from a denticle was 32.57 ug/ul (Fig 1). When compared to concentrations of DNA extracted from tissue we see no significant difference (f=3.56, df=2,10, p> 0.05). The purity of the extractions averaged 1.97 260/280λ for fragments and 1.38260/280λ for denticles (Fig 2). We saw no significant difference in purity due to origin of the DNA (t=-2.58, df=1.067, p>0.05).

**Fig 1.**
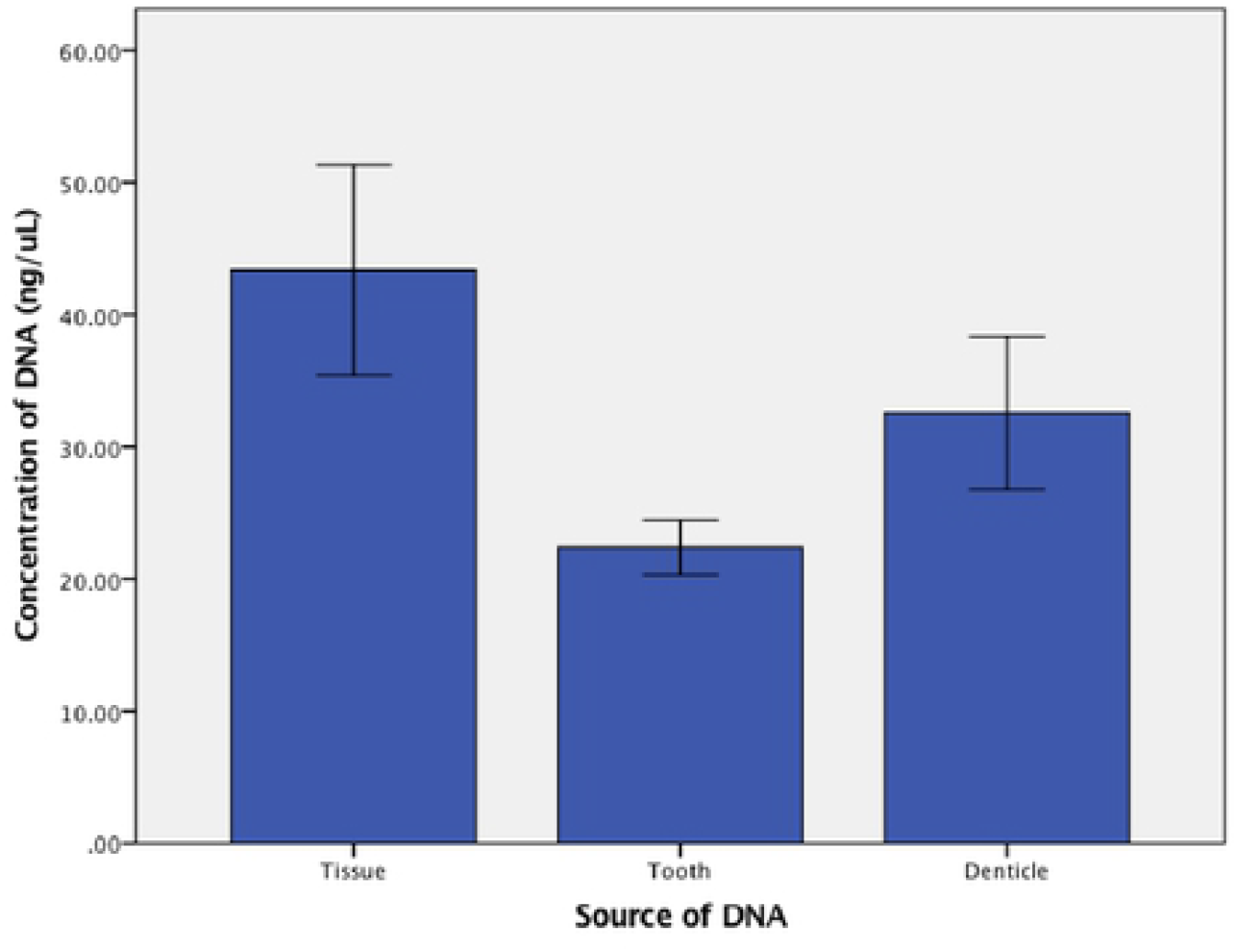
DNA concentration comparison. Comparison of DNA concentrations from Tissue, Tooth and Denticle Enamel. Error bars represent standard error of the mean.

**Fig 2.**
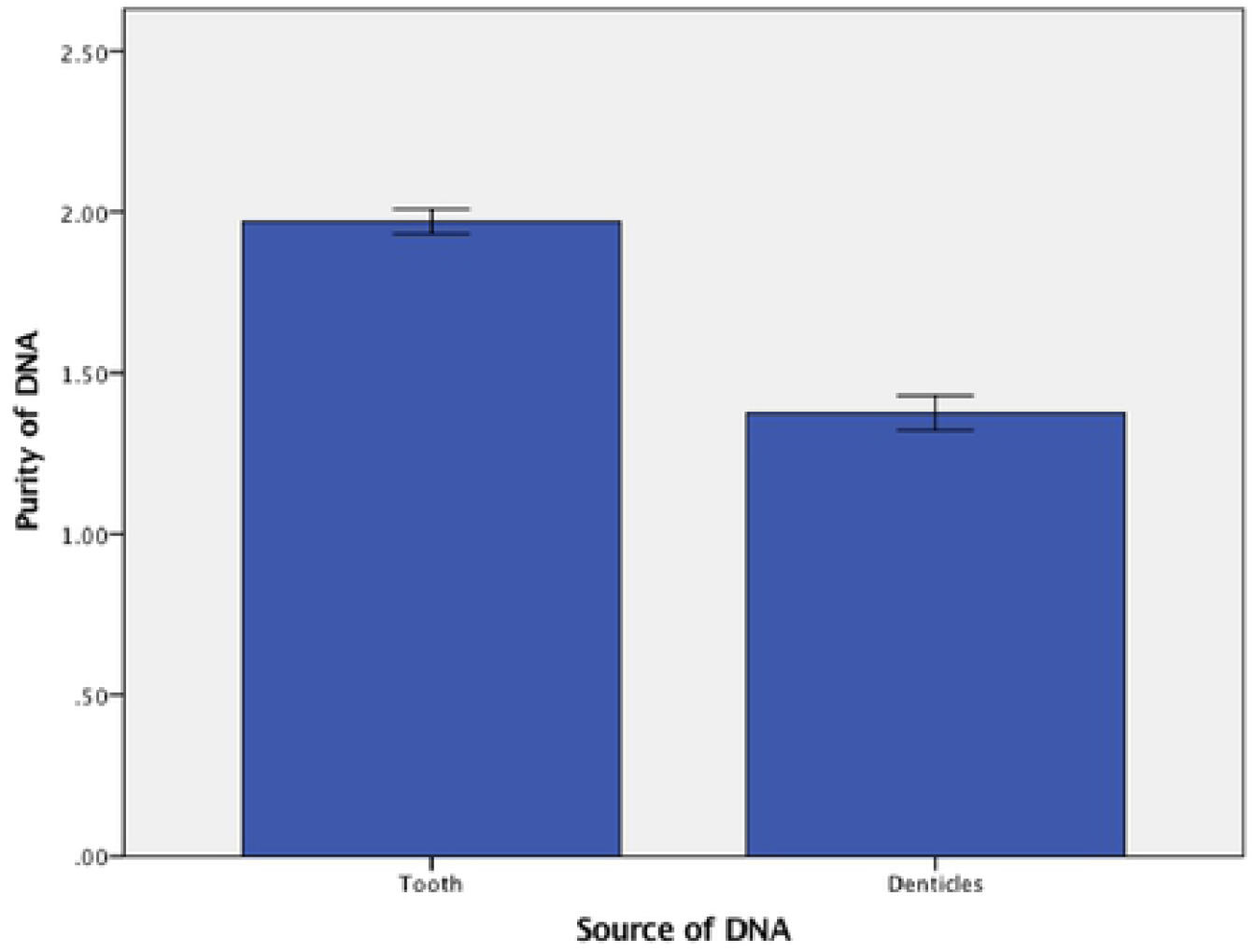
DNA purity comparison. Comparison of DNA purity from Denticle and Tooth Enamel. Error bars represent standard error of the mean.

The high mineral concentration in the teeth and denticles also interferes with DNA amplification. Decalcification helps address this latter problem, improving DNA amplification success of all extracts. DNA did not seem to uniformly distributed in all layers of the tooth or denticle for this reason, we utilized 2–3 extracts from each sample to address the seemingly random distribution of DNA. Using multiple extracts enables one to determine if the sample has sufficient DNA to warrant further processing. If all extracts fail to yield a product PCR amplification of the extracted DNA, is not suggested, unless it has considerable biological uniqueness.

This technique of extraction could allow for the possibility of new noninvasive collection techniques to be used in the sampling of Lamniforms. Furthermore, this methodology could allow for the comparison of historic samples/ fragments from attacks to those of traditionally collected samples.

## Acknowledgements

We would like to thank the following individuals for their assistance in this project; Keith Poe, Dirk Schmidt, Sam Wasser, Tori Thompson, Elena Shink and Hayley Verner.

